# Proteins encoded by Novel ORFs have increased disorder but can be biochemically regulated and harbour deleterious mutations

**DOI:** 10.1101/562835

**Authors:** N. Suhas Jagannathan, Narendra Meena, Kethaki Prathivadi Bhayankaram, Sudhakaran Prabakaran

## Abstract

Recent advances in proteogenomics indicate that protein or protein-like products can be encoded by previously uncharacterized Open Reading Frames (ORFs) that we define as Novel Open Reading Frames (nORFs)^1,2^. Although it is yet unclear if these protein or protein-like products could possess any significant biological function hopes have been raised to target them for anticancer and antimicrobial therapy ^3,4^. In this study, we used computational tools to systematically investigate these novel protein sequences for their propensities toward structural disorder, post-translational modifications (PTM) and mutational densities. We found that these novel proteins have significantly higher disorder and similar PTM frequencies compared to known proteins. Although these regions were found to harbour deleterious mutations, we did not observe any correlation between the pathogenicity of mutations and their location (ordered/disordered) within these novel proteins. This study suggests that these nORFs encode an important class of proteins, that could undergo sequence, structural or regulatory changes during complex diseases, and hence warrant further study.

**Significance:** Certain noncoding regions of the genome are known to make protein-like products. These protein-like products have significantly expanded the number of the cellular proteome but we dont know whether they are capable of performing biological functions. Here in this study, we have investigated all such putative protein-like products and analyzed whether they can form structures, whether hey harbour mutations, and whether they can be regulated by biochemical regulatory processes. Results from this study indicate that indeed these protein-like proteins can perform and be involved in all the above processes.

---

Over 500 million years of evolution from hydra to humans, the total number of ORFs have been thought to remain the same at around 30,000. With the advent of deep sequencing strategies in both genomics and proteomics fields, we are now discovering nORFs that have remained undiscovered or ‘hidden’ ^1,2,5,6^. These nORFs are pervasive throughout the genome and are observed in both the coding and non coding regions ^1,7^. They are variously classified as small ORFs (sORFs) ^8,9^ which are 1-100 amino acids in length, altORFs ^10^, which are proteins in alternate frames to known proteins, Denovogenes^11^ or Orphan genes ^12^, Pseudogenes ^1,13^, and many ncRNAs have been shown to have coding potential ^14–18^. These new discoveries challenge traditionally held conservative definitions of ORF, as used until the recent past ^19^.

While these new discoveries have illuminated that the cellular proteome is much more complex than our current understanding, there is a huge knowledge gap on the putative functions of nORFs. There have been two lines of speculations about it: on one side some have dismissed the novel proteins as mere biological noise ^20^, while on the other side some have proposed that such nORFs confer evolutionary advantage ^21,22,23^ to an organism. Without a systematic structural investigation of these novel proteins, it is difficult to ascertain which of these two hypotheses might be true.

To investigate these conflicting hypotheses, we first curated a list of all nORFs that have been identified with evidence of translation. We obtained sequences for known and verified human proteins from NeXtProt (https://www.nextprot.org/) ^24^, sequences for sORFs from the sORF database (http://sorfs.org/database) ^8^, sequences for altORFs from Roucou’s lab ^10^ (http://haltorf.roucoulab.com/), and sequences of Pseudogenes with evidence of translation from Xu et al ^13^. For Denovogenes, we manually curated a list of 42 protein sequences through literature search. For conservative measurements of disorder scores, we discarded protein sequences less than 30 amino acids in length from all these datasets, since these were likely to be enriched for disorder. Non coding RNA sequences were downloaded from RNACentral database (http://rnacentral.org) ^25^. While all the other datasets contained protein sequences whose translation has been experimentally verified in literature, the downloaded RNA central dataset contained 9,386,637 nucleotide transcript sequences. We identified potential ORFs from these transcripts, using the following workflow. Each sequence was subjected to three-frame translation using the EMBOSS *transeq* program provided as a standalone utility by EMBL-EBI. From the output protein sequences, putative translated ORFs were obtained by identifying all possible subsequences (>30 residues in length) beginning with a Methionine and ending at a STOP codon (EMBOSS *checktrans* program and Matlab scripts to parse the output text files). After removing redundant sequences from the extracted list, we obtained a unique set of 5,185,186 protein sequences, which we used as putative transcripts from the RNAcentral database for disorder prediction. Since the size of the RNACentral dataset far exceeded that of the four other novel datasets, we decided to keep the datasets segregated for future analysis. From here on we will call the amino acid sequences encoded by the nORFs as novel proteins. **Figure 1** shows the protein length distributions for each of the studied datasets. We found that the novel proteins are shorter in length compared to known proteins. Next, we investigated the structure-forming capabilities of these novel proteins, since shorter proteins are known to form elementary structures ^26^. We employed two disorder prediction algorithms, PONDR (*http://www.pondr.com*) and IUPred (*https://iupred2a.elte.hu/*), to assess whether these novel proteins are predominantly ordered or disordered, which would directly correlate with their ability to form structures.

**Figure 1.**
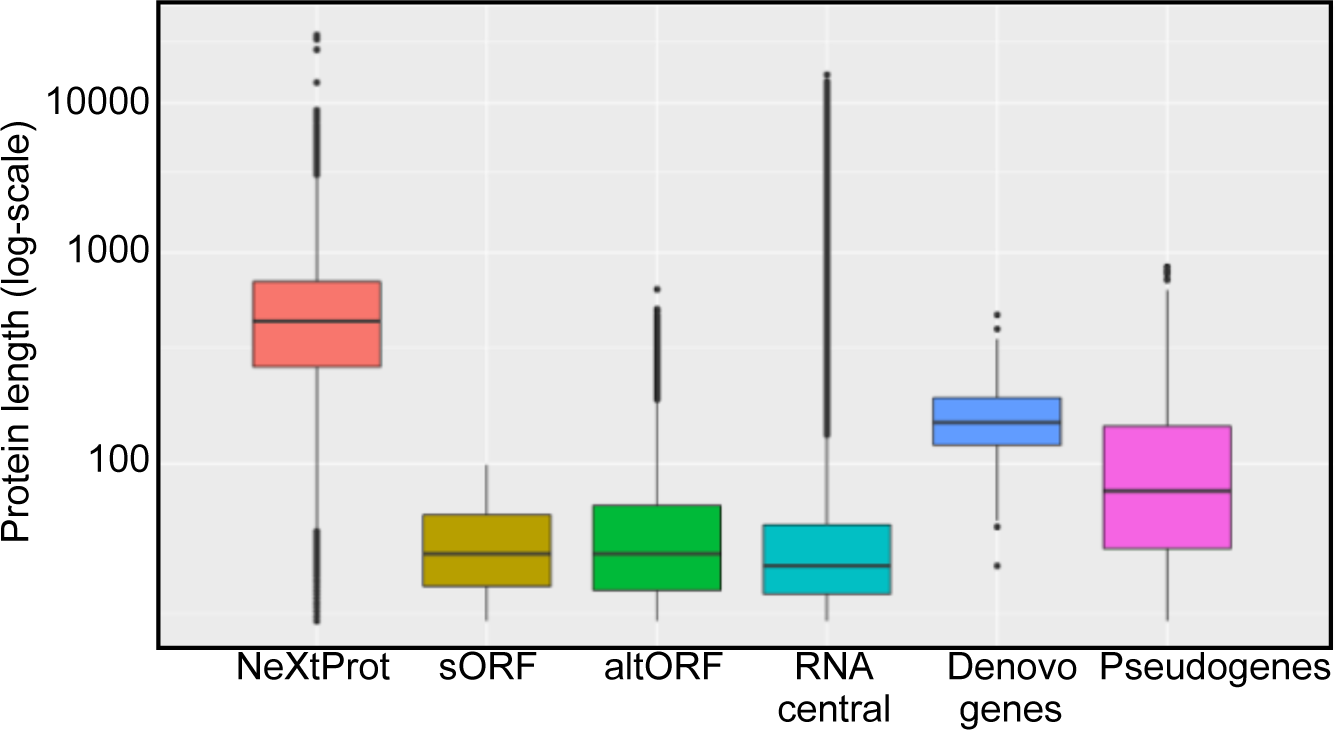
Amino acid length distribution of novel proteins encoded by nORFs. Amino acid sequences of known human proteins from NeXtProt, and potential proteins encoded by nORFs - sORFs, altORFs, Pseudogenes, Denovogenes, and all possible translated amino acids sequences from RNA central, were retrieved as described in the main text. Shown are the distributions of protein length in each dataset. The nORFs on average have shorter sequences compared to the structured proteins in the NeXTProt database.

PONDR is an algorithm that uses feedforward neural networks and sequence attributes of amino acid windows (9-21 AAs) to predict disorder. Among PONDR-based algorithms, we used the VSL2 algorithm that was optimized and trained using both short and long protein sequences. IUPred identifies Intrinsically Disordered Protein Regions (IDPRs, i.e. regions that lack a stable monomeric structure under native conditions) based on a biophysics-based model. Among the three IUPred-based algorithms, we performed separate predictions with IUPred ‘long disorder’ and IUPred ‘short disorder’. Matlab-based scripts were written to automate and batch process protein sequences for disorder prediction, and parse the output. For both PONDR and IUPred, the output consisted of average disorder score (in the range 0-1) (**Fig. 2A-C**) for a protein sequence, and the percentage of each sequence that was predicted to be disordered (**Supplementary Fig. 1A-C**). A protein sequence was considered “disordered” when the predicted average disorder score was greater than 0.5. The computed bootstrap confidence intervals of mean (and median) average disorder scores showed that the nORF datasets (sORFs, altORFs, RNACentral, Pseudogenes and Denovogenes) had higher mean (and median) values of disorder, compared to known proteins in NeXtProt (**Fig. 2D**). We also checked to see if these novel protein datasets were enriched for disordered proteins using Fisher’s exact test and the Chi-square test. Our tests showed that each of the nORF groups (except Denovogenes) was enriched for disordered sequences in comparison to proteins in NeXtProt (**Fig. 2E**). Supplementary Fig. 1D shows the final number of amino sequences in each novel protein category used in the above analysis. All statistical tests were corrected for multiple hypothesis testing, using FDR values computed by the Benjamini Hochberg method.

**Figure 2.**
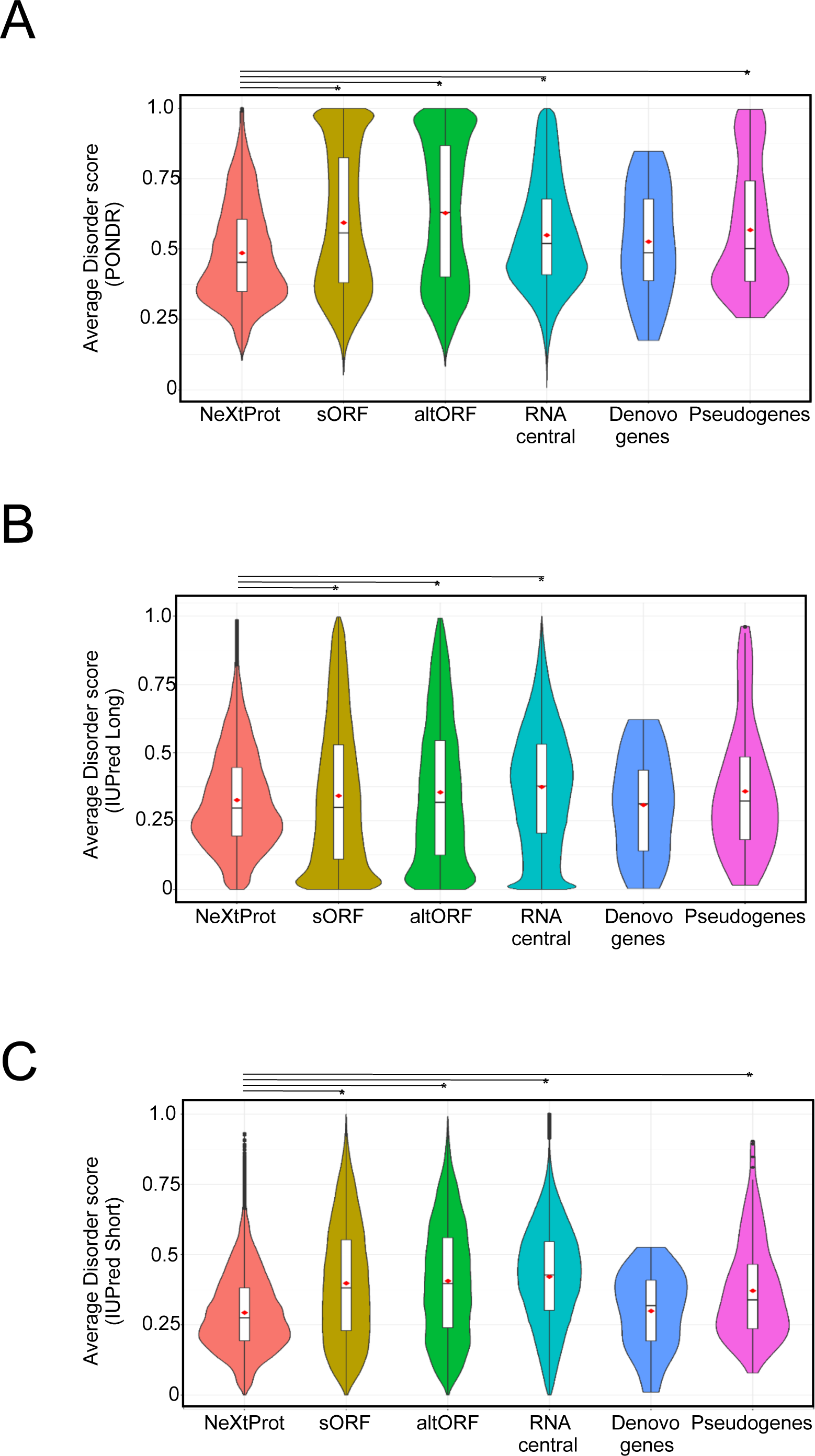
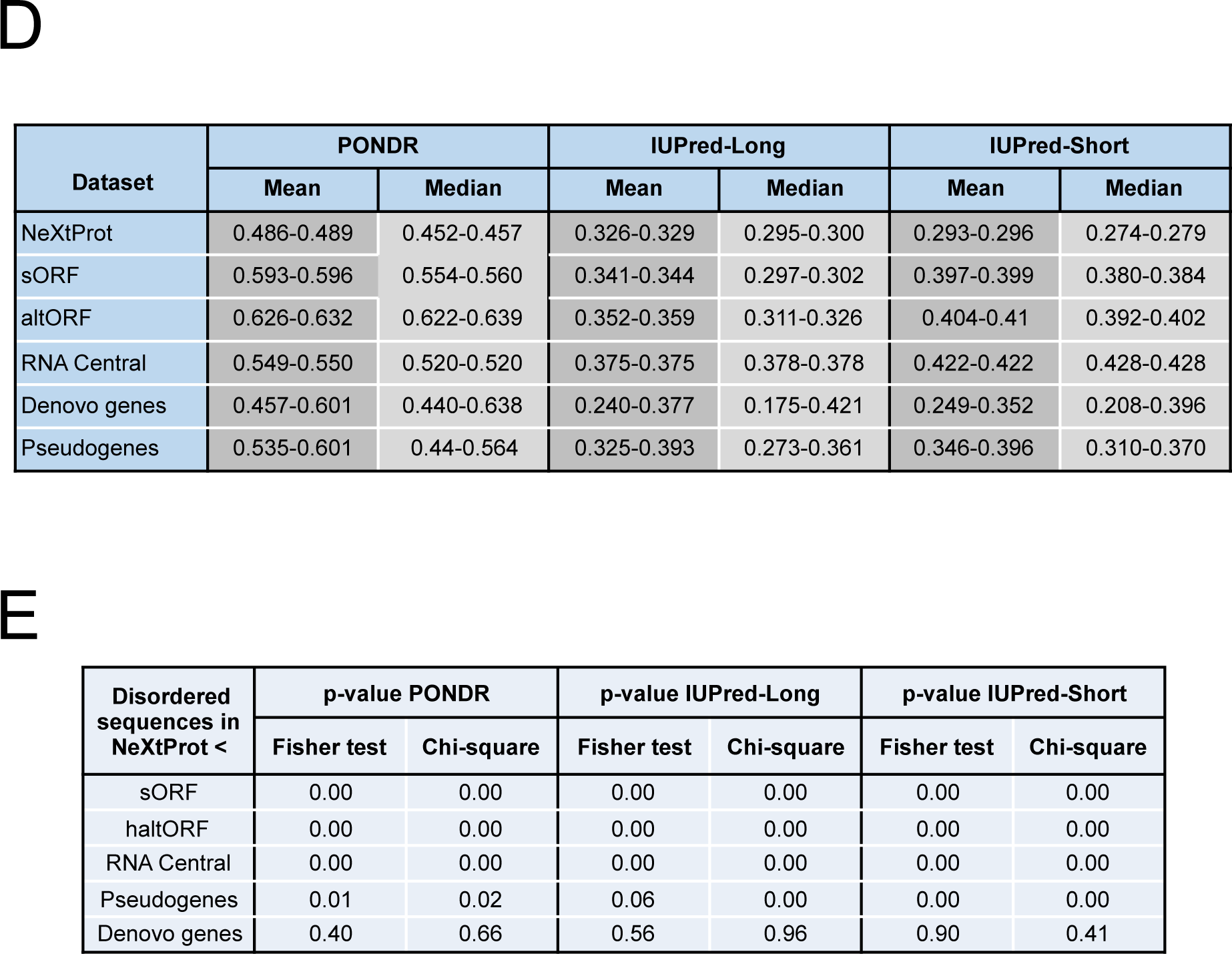
Prediction of disorder in proteins encoded by nORFs. Average disorder scores of proteins in NeXtProt compared to average disorder scores of proteins encoded by nORFs, predicted by **A.** PONDR, **B**. IUPred-Long, and **C.** IUPred-Short disorder predictors. **D.** 95% Bootstrap confidence Interval of the mean and median of disorder scores predicted using PONDR, IUPred-Long and IUPred-Short for each of the nORF datasets **E.** Statistical significance (uncorrected p-values) for the enrichment of disordered sequences in nORF datasets, in comparison to NeXtProt.

Although the average disorder score and the percentage disorder scores are higher in nORFs, they are not drastically high to completely negate or rule out the structure forming capability of novel proteins. Some disordered regions have been known to undergo disorder-to-order transitions upon binding to substrates. We used the Anchor program (*http://anchor.enzim.hu*) ^27^ to investigate whether the novel proteins show increased propensity to form structures. The results of this analysis indicate that novel proteins, except for Denovogenes, show increased anchor scores compared to NeXtProt proteins (**Figure 3A**). However, we also found a strong positive correlation between average anchor score and average disorder score for most data sets, which is not surprising, since the prediction of binding sites uses biophysical parameters similar to those involved in disorder prediction (**Figure 3B**).

**Figure 3.**
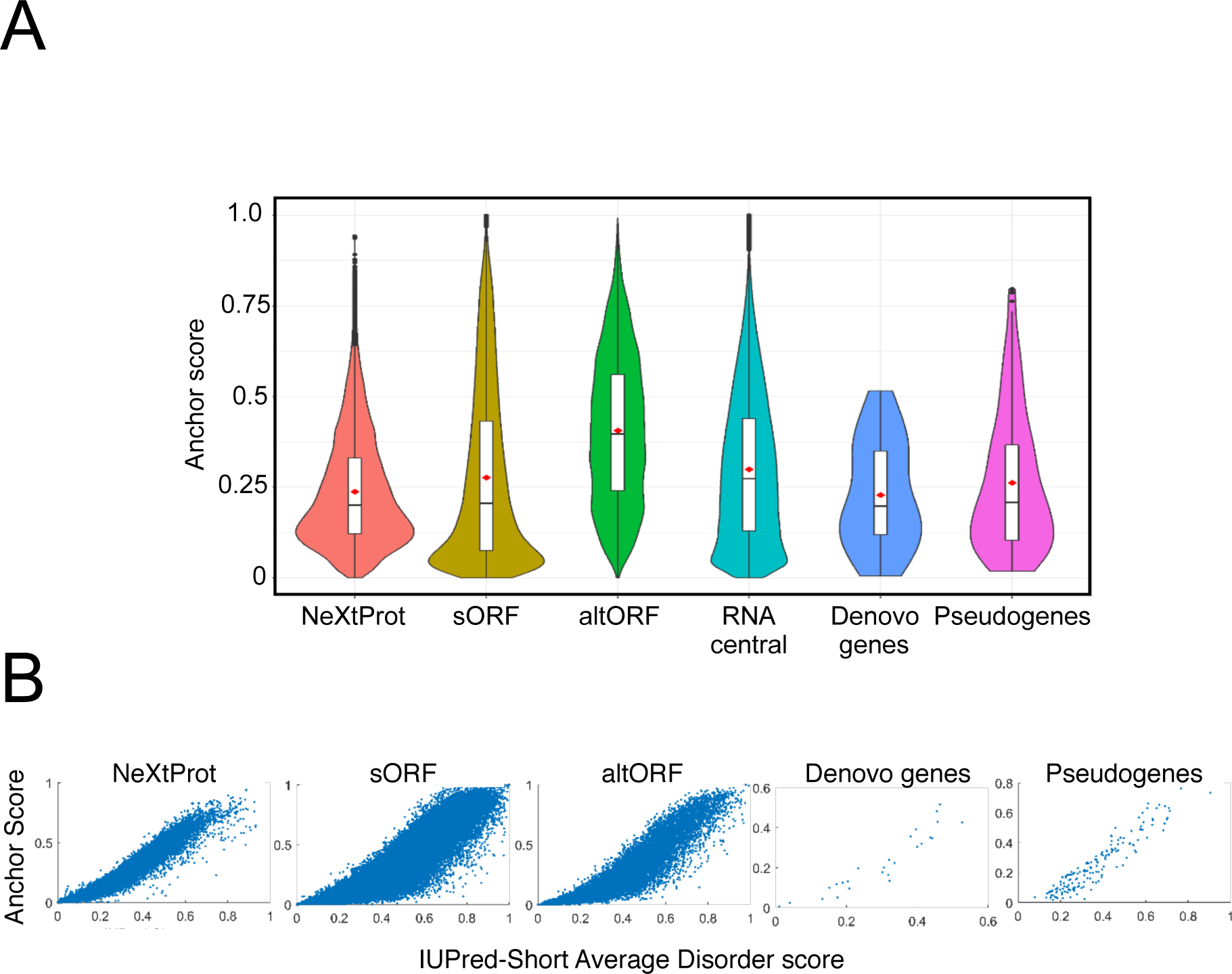
Anchor predictions of binding regions in proteins encoded by nORFs. Anchor scores represents the average propensity of an amino acid in a disordered region to be part of a protein-protein binding site. **A.** Shown are the distributions of the predicted Anchor scores for known proteins in NeXtProt, and those encoded by the nORF datasets. As expected, the nORF proteins have higher mean Anchor scores in comparison to the NeXTProt database. **B.** Observed correlation between average Anchor score and IUPred-Short disorder prediction score for individual datasets.

Given that the novel proteins were found to be enriched for disorder, we investigated whether they might be biologically regulated by biochemical mechanisms such as Post Translational Modifications (PTM). Previous evidence suggested that novel proteins are indeed biologically regulated ^1^ and they may be enriched for regulatory sites, but the observation was not done at a global scale. Hence, we predicted possible PTM sites in the amino acid sequences from all novel proteins, using the ModPred stand-alone software ^28^. For each sequence, we predicted amino acid sites for nine PTMs - Phosphorylation, Acetylation, Methylation, Sulfation, SUMOylation, Ubiquitination, C-linked, O-linked and N-linked glycosylation (**Figure 4**)

**Figure 4.**
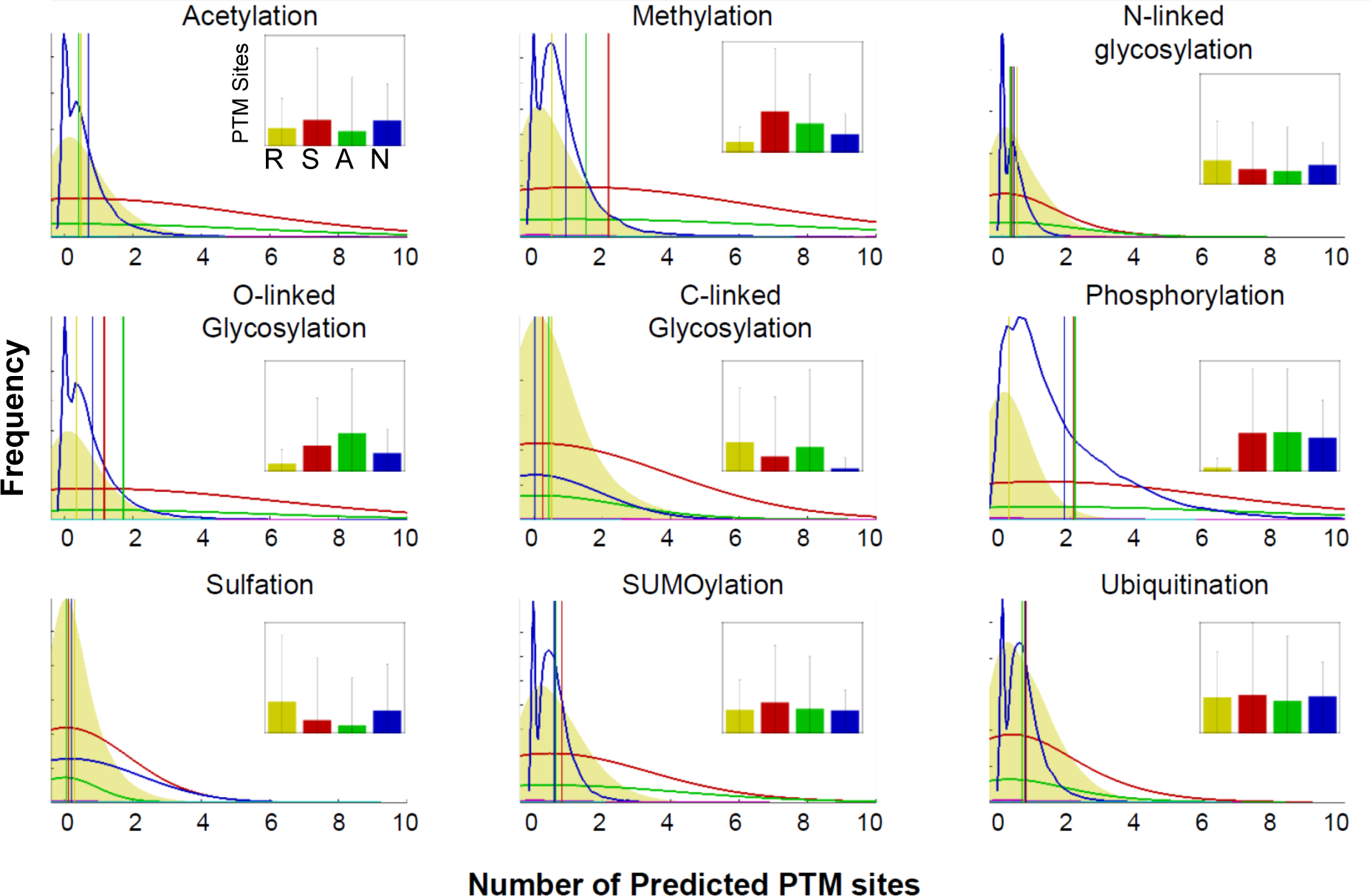
Predictions of PTM site-densities in the proteins encoded by nORFs. PTM sites in protein sequences were predicted using the ModPred tool. Predicted densities of nine PTM modifications for sORFs, altORFs, and NeXtProt sequences were compared with the predicted PTM densities for a pseudo-dataset composed of 500,000 randomly generated amino acid sequences (100-aa long). In each panel, the shaded yellow histogram represents the predicted distribution of the number of PTM sites in the random dataset. The curves (red and green, blue) represent the predicted density distributions for sORF, altORF, and NeXtProt respectively. The overlaid vertical lines show the mean of each distribution. For many cases (e.g., Methylation), it can be clearly seen that the average number of predicted PTMs is higher in all three datasets compared to the random set (all three lines are to the right of yellow). Shown in insets are bar plots of the same data indicating the PTM density distributions of the random data set (R, yellow bar) vs sORF (S, red), altORF (A, green) or NeXtProt (N, blue).

To compare if each of the datasets have higher or lower predicted PTM site-densities than expected at random, we generated 500,000 random protein sequences of 100 amino acids each, from a uniform distribution of 20 amino acid propensities. ModPred was then used to predict PTM sites in these random sequences for the same list of nine modifications. The number of predicted PTM sites in all datasets were normalized to account for variable sequence length. We compared the distribution of predicted PTMs in each of the nORF datasets, and NeXtProt against the predicted PTM distributions corresponding to the dataset of random sequences. Methylation, O-linked glycosylation, and Phosphorylation were found to be significantly enriched in both novel proteins and known NeXtProt proteins, compared to the random dataset (p < 0.0001, Wilcoxon rank sum test). Overall, for most PTM types, the densities of predicted PTMs was comparable, if not higher, in the novel proteins versus the NeXtProt database (**Figure 4**). This indicates that the novel proteins could be subjected to any biochemical regulation just as much as all known proteins.

To investigate whether the novel protein regions could harbour disease-associated mutations, we mapped mutations from the Catalogue of Somatic Mutations in Cancer (COSMIC) and Human Gene Mutation Database (HGMD) databases to the novel proteins. **Figure 5A-D** shows examples of COSMIC or HGMD mutations mapped to sORFs, Denovogens, and Pseudogenes demonstrating that these regions do indeed harbour mutations. Finally, we investigated whether the pathogenicity scores of these mutations, assessed as Combined Annotation Dependent Depletion (CADD) ^29^ and Functional Analysis through Hidden Markov Models (FATHMM) ^30^ scores, had any correlation with disorder scores at the mutated region of the novel proteins (both amino-acid specific disorder score, and average disorder score for a 7-aa window around the mutated residue). This analysis (**Figure 6A-D** and Supplementary Fig. 2) did not reveal any correlation between low pathogenicity and higher disorder scores.

**Figure 5.**
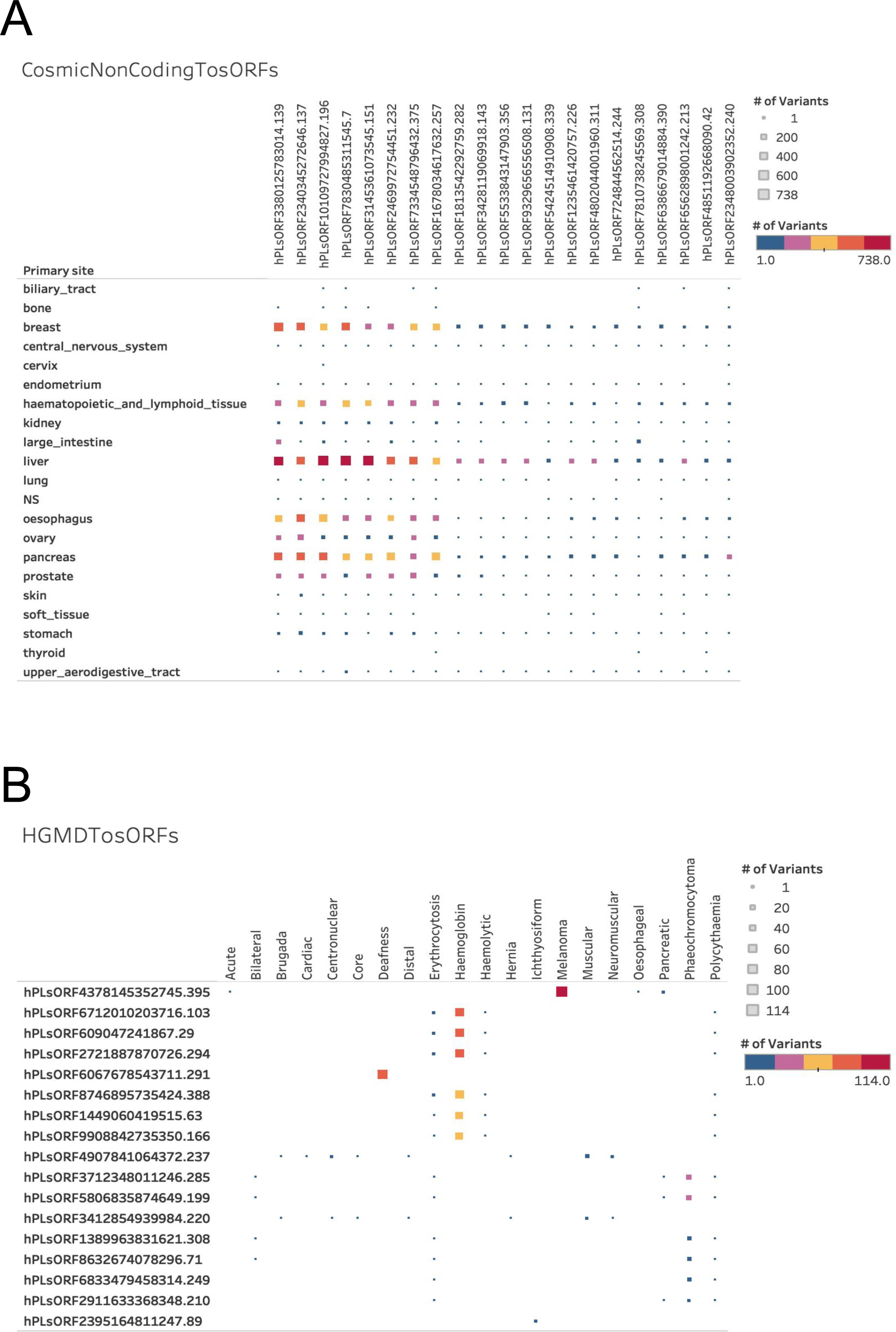
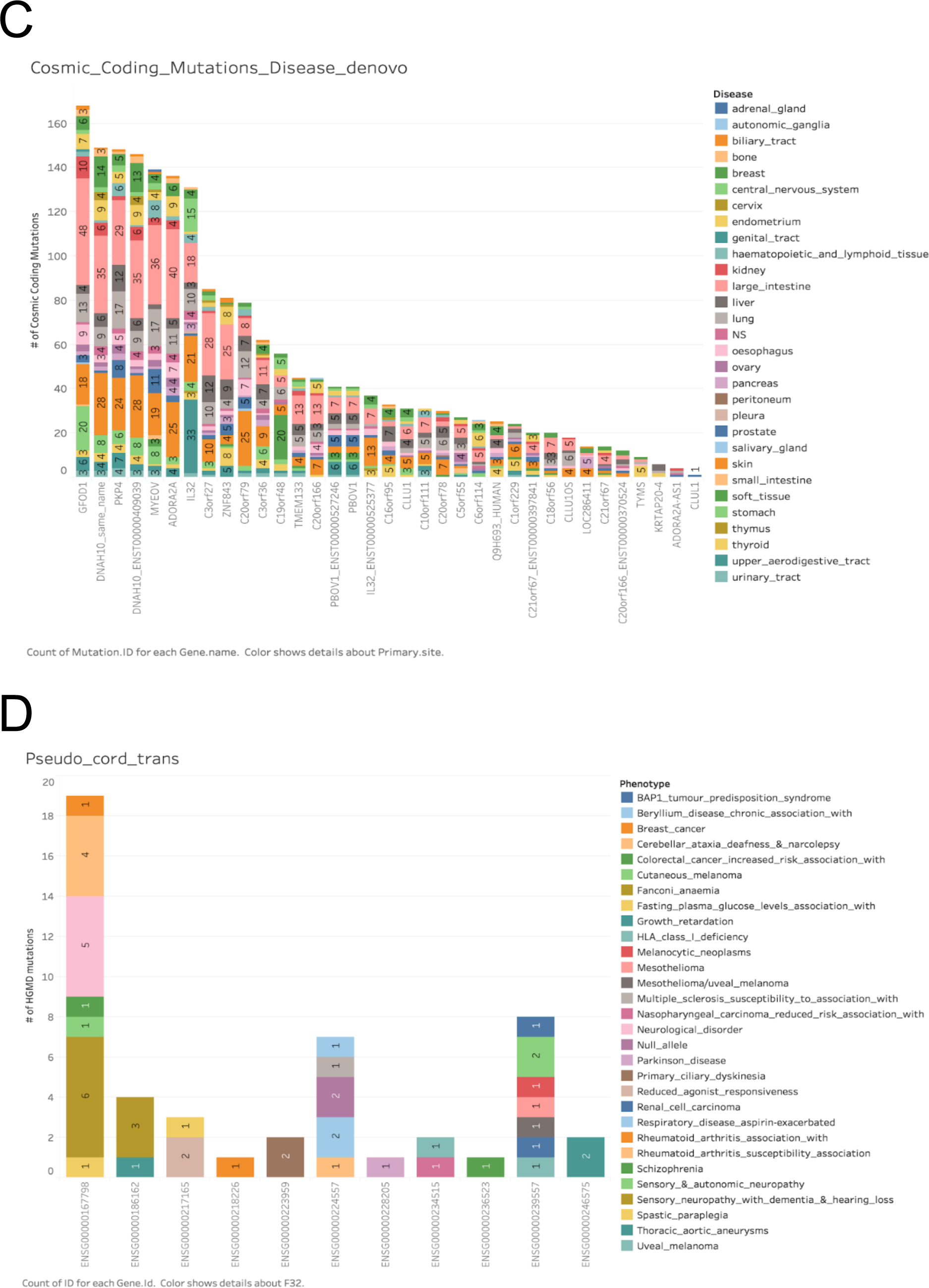
Known mutations in the proteins encoded by the nORFs. Mutations from COSMIC and HGMD databases were mapped to both entire nORFs genomic and specifically to novel protein amino acid sequence coordinates and represented as described below. **A.** Noncoding mutations from COSMIC were mapped to all sORFs genomic coordinates and only the top 21 sORFs with the highest number of mutations are represented here. Size of the squares against each sORF and each disease indicates the number of mutations and the color indicates the number of pathogenic mutations in that region. **B**. Mutations from HGMD database were mapped to sORFs and only the top 17 mutations are represented here. **C.** Coding mutations from COSMIC are mapped to all Denovo genes and are represented. **D.** Mutations from HGMD are mapped to pseudogenes that are known to be translated and presented.

**Figure 6.**
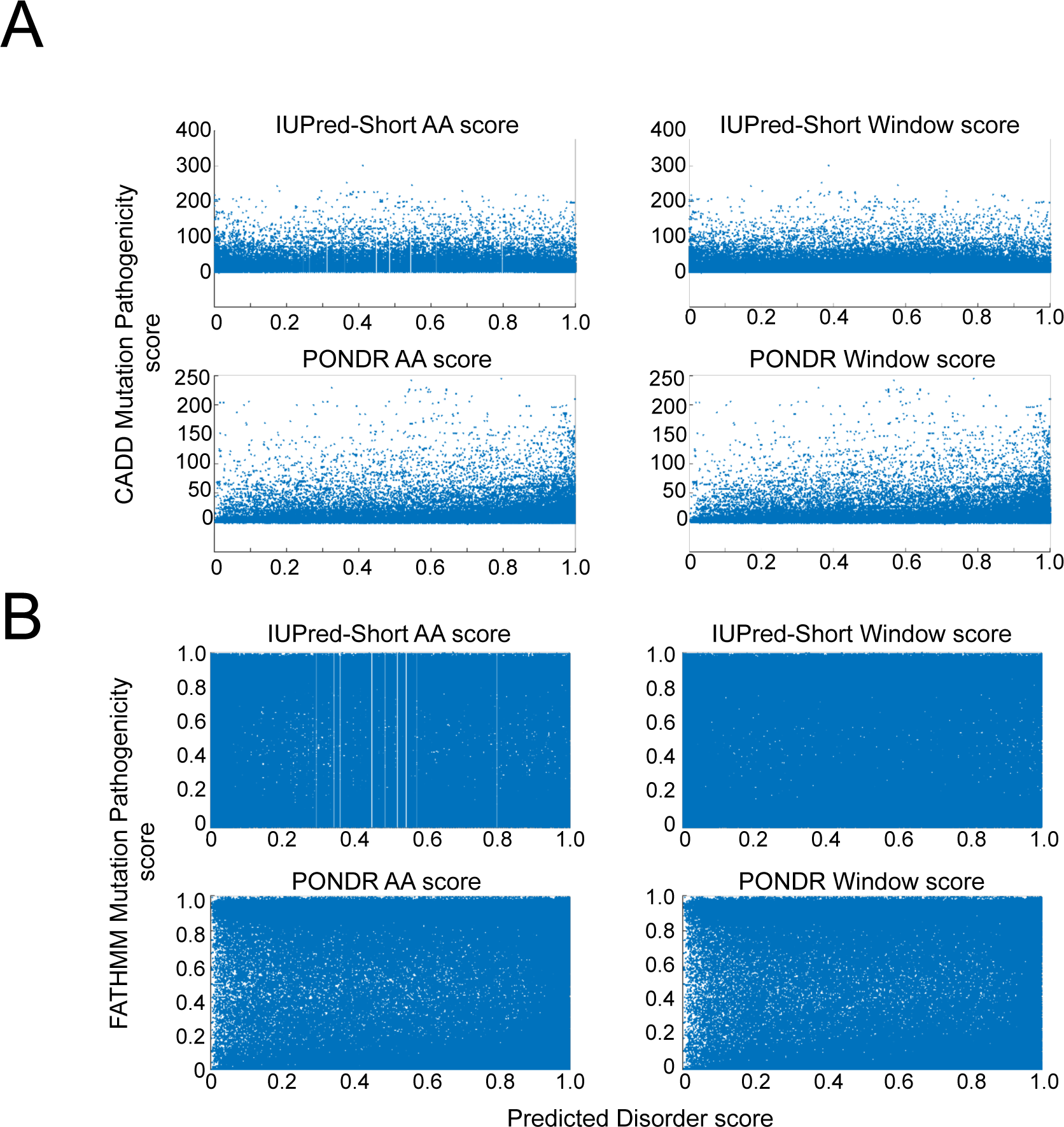
CADD and FATHMM pathogenicity scores vs predicted disorder scores for proteins encoded by sORFs. We plotted the CADD or FATHMM mutation pathogenicity scores for the proteins encoded by sORFs, against their corresponding disorder scores predicted using either PONDR and IUPred. Disorder scores were computed at either amino-acid resolution, or for a 7-AA window around the mutated residue. The analysis did not reveal any correlations between **A.** FATHMM scores and predicted disorder scores, or **B.** CADD scores and predicted disorder scores.

Taken together our results demonstrate that although the novel proteins have increased disorder the proportion of increase is not substantial to affect their structure forming capabilities. In addition they can be biochemically regulated by PTMs and more importantly harbour deleterious mutations that is not correlated with whether those regions are ordered or disordered, suggesting that the novel proteins could be involved in biochemical processes in cells that may be dysregulated in diseases. Hence we should investigate the functional role these novel proteins and nORFs in depth.

## Acknowledgements

We would like to thank Asst Prof. Lisa Tucker-Kellogg (Duke-NUS Medical School, Singapore) for kindly supporting NSJ to work on this project

## Author Contributions

NSJ performed most of the analysis, participated in writing the manuscript. NM performed the mutational analysis. KB did the Denovogenes and Pseudogenes mutational analysis. SP designed and supervised the project, analysed the data, and wrote the manuscript; Competing interests: SP is a co founders of NonExomics; and Data and materials availability: Almost all processed data is in the main text or in the supplementary materials.

## SUPPLEMENTARY INFO

**Figure S1 A-C.**
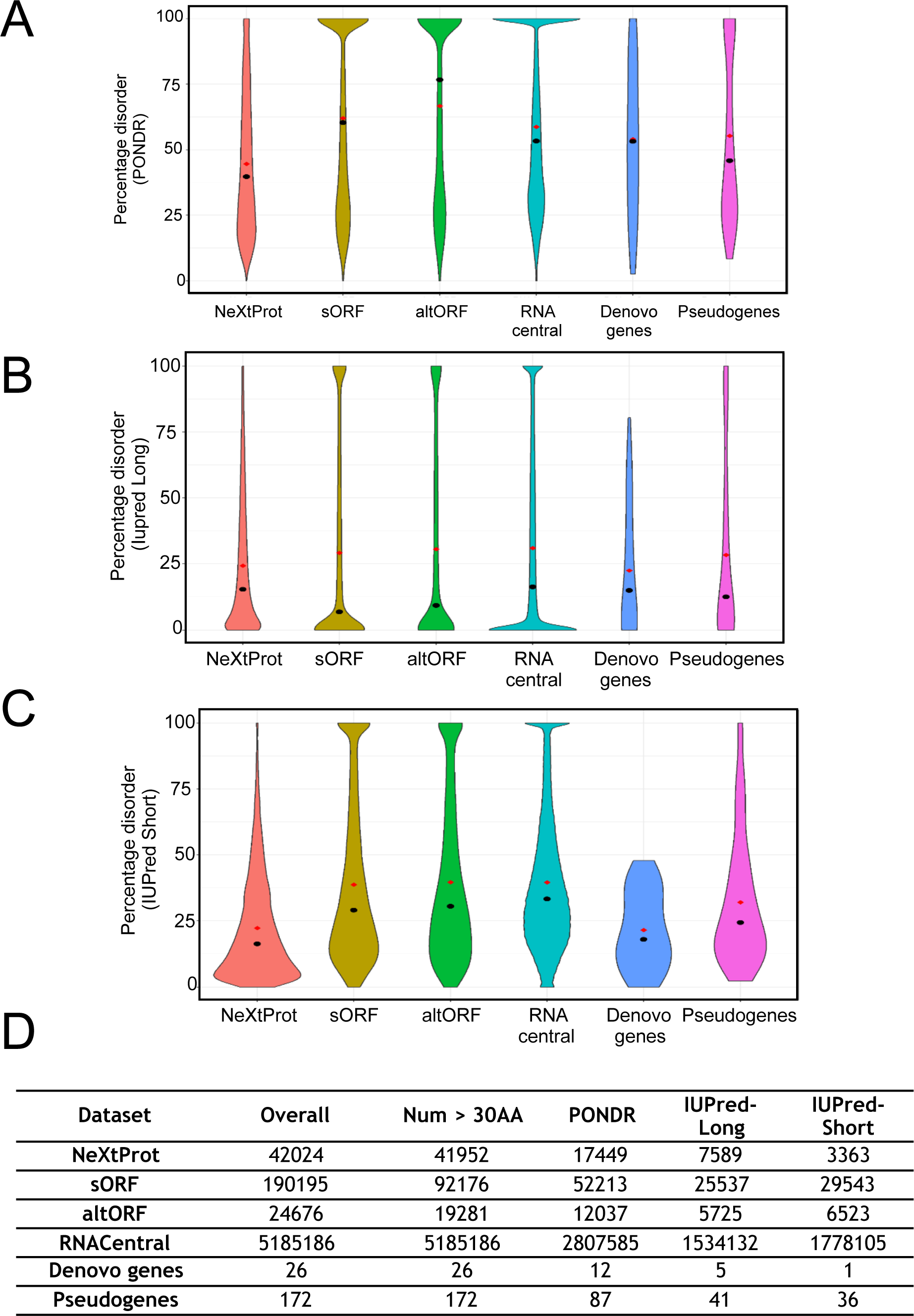
Percentage of protein sequence identified to be disordered (amino-acid disorder score > 0.5) in NeXtProt, and each of the nORF datasets, for three prediction algorithms PONDR, IUPred-Long and IUPred-Short. Shown in red and black within each distribution are the mean and median respectively. **D.** Table showing the number of protein sequences identified in each nORF dataset, number of sequences used for further analysis (Sequence length > 30) and the number of predicted disordered sequences (average disorder score > 0.5) obtained using PONDR, IUPred-Long and IUPred-short algorithms.

**Figure S2.**
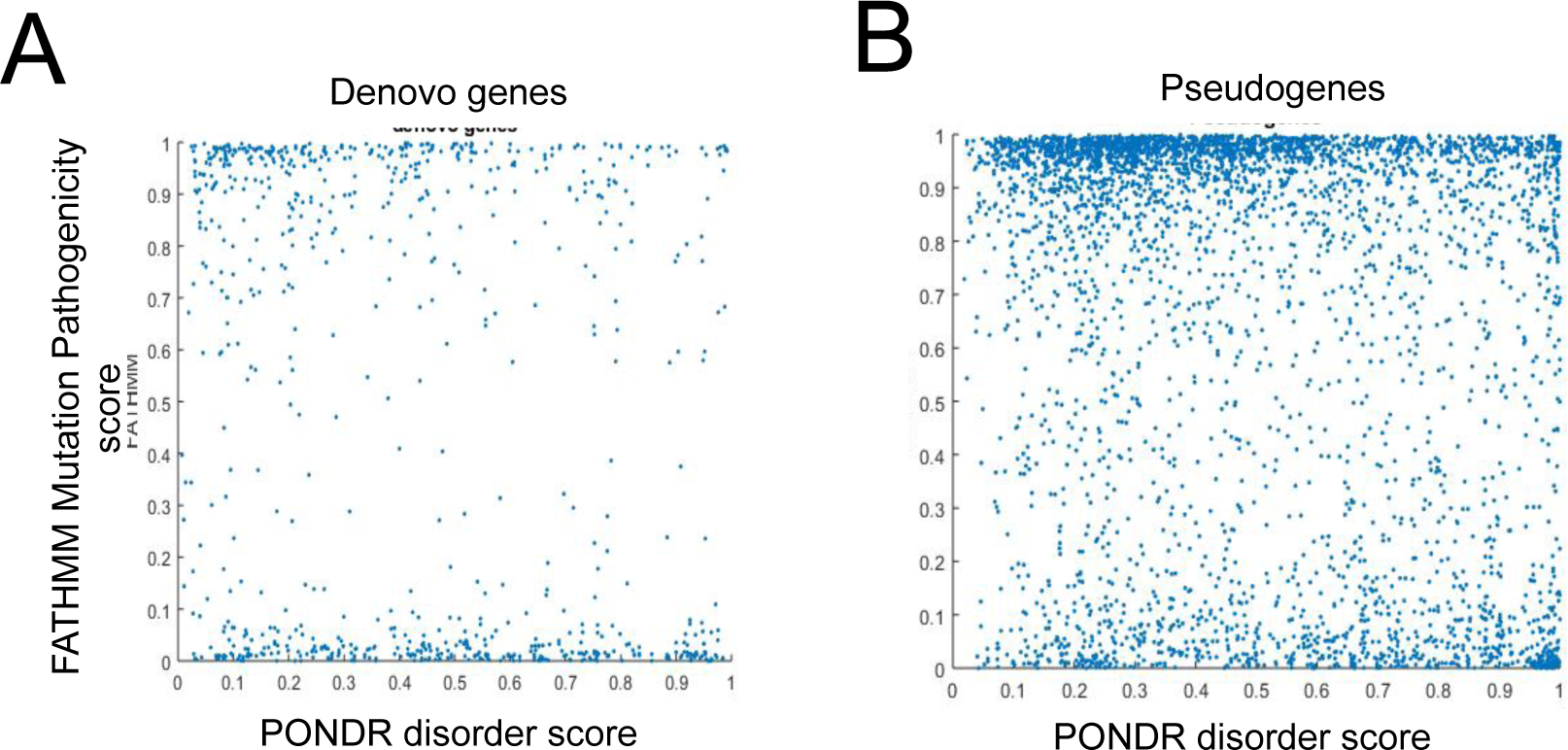
Computed FATHMM scores vs PONDR-predicted disorder scores for **A.** Denovo genes or **B.** Pseudogenes. No correlation between mutation pathogenicity and disorder score was observed in either case.

